# Modelling Instantaneous Firing Rate of DBS Target Neuronal Ensembles in Basal Ganglia and Thalamus

**DOI:** 10.1101/2022.06.28.497834

**Authors:** Yupeng Tian, Matthew J. H. Murphy, Leon A Steiner, Suneil K Kalia, Mojgan Hodaie, Andres M Lozano, William D Hutchison, Popovic Milos R., Milosevic Luka, Lankarany Milad

## Abstract

**Objective:** Deep Brain Stimulation (DBS) is an effective treatment for movement disorders, including Parkinson’s disease and essential tremor. However, the underlying mechanisms of DBS remain elusive. Despite the capability of existing models in interpreting experimental data qualitatively, there are very few unified computational models that quantitatively capture the dynamics of the neuronal activity of varying stimulated nuclei—including subthalamic nucleus (STN), substantia nigra pars reticulata (SNr) and ventral intermediate nucleus (Vim)—across different DBS frequencies.

**Materials and Methods:** Both synthetic and experimental data were utilized in the model fitting; the synthetic data were generated by an established spiking neuron model that was reported in our previous work ^1^, and the experimental data were provided using single-unit microelectrode recordings (MER) during DBS (microelectrode stimulation). Based on these data, we developed a novel mathematical model to represent the firing rate of neurons receiving DBS, including neurons in STN, SNr and Vim—across different DBS frequencies. In our model, the DBS pulses were filtered through a synapse model and a nonlinear transfer function to formulate the firing rate variability. For each DBS-targeted nucleus, we fitted a single set of optimal model parameters consistent across varying DBS frequencies.

**Results:** Our model accurately reproduced the firing rates observed and calculated from both synthetic and experimental data. The optimal model parameters were consistent across different DBS frequencies.

**Conclusion:** The result of our model fitting were in agreement with experimental single-unit microelectrode recording data during DBS. Reproducing neuronal firing rates of different nuclei of the basal ganglia and thalamus during DBS can be helpful to further understand the mechanisms of DBS, and to potentially optimize stimulation parameters based on their actual effects on neuronal activity.

## INTRODUCTION

Parkinson’s disease (PD) is a common neurodegenerative disorder characterized by motor symptoms like stooped posture, shuffling gait (festination), akinesia, rigidity, and rest tremor ^2, 3^. Essential tremor (ET) is a common movement disorder affecting up to 1% of adults over 40 years of age, and is featured by attention tremor and uncontrollable shaking of the affected body part ^4, 5^. Development of PD and ET symptoms is thought to be related to the pathological changes in the basal ganglia, thalamus and cerebellum ^1, 4^. Deep brain stimulation (DBS) has become a standard therapy for movement disorders, including PD ^6^, ET ^7^, and dystonia ^8^. DBS has also been investigated for therapeutic effects of psychiatric and cognitive disorders, including obsessive-compulsive disorder ^9^, Alzheimer’s disease ^10^ and epilepsy ^11^. Despite the established benefits of DBS, its therapeutic mechanisms on neuronal activity are not fully understood ^1, 12^. Synaptic depression, which can stem from synaptic and axonal failure ^13^, was suggested as the main biophysical explanation to the intermittent firing patterns of stimulated nucleus in high frequency DBS ^14^. Recently, Milosevic et al. ^1^ demonstrated that different frequencies of DBS modulate the firing rate of the stimulated nucleus differently. During low frequency DBS, the ratio of excitatory and inhibitory pre-synaptic inputs influences the firing rate of stimulated neurons. In high frequency DBS, stimulated neurons are often suppressed due to synaptic depression ^1^. For STN neurons, during high frequency DBS, the excitatory pre-synaptic inputs are suppressed ^15^, whereas the inhibitory pre-synaptic inputs persist due to the increasing activation of GABAB receptors ^16, 17^. Nevertheless, detailed dynamics underpinning the experimental firing rate of the stimulated neurons in response to different DBS frequencies were left unknown due to the lack of biophysically-reasonable models that allow to link those dynamics to observed firing rates. These models enable predicting changes of neuronal activities of stimulated neurons.

The firing rate of a population of neurons—which might be modulated by DBS—is a representative feature of the underlying neuronal dynamics, and has been widely used in the modeling of the sensory cortex ^18, 19^, the visual cortex ^20, 21^, Parkinson’s Disease ^22^ and cultured network ^23^. Despite the significance of firing rate, the existing models on DBS-induced neuronal dynamics are often based on the variability in the membrane potential, or oscillations observed in the local field potentials (LFP) ^1, 4, 24^. The spiking neuron models aim to replicate the membrane potential dynamics of single neurons ^1, 13^, while abstract models were developed to track neuronal activities recorded by macro electrodes, e.g., LFP ^4, 25^. Milosevic et al. ^1^ utilized a model for short-term synaptic plasticity, together with a leaky integrate-and-fire (LIF) model, to track firing patterns of stimulated nuclei qualitatively. Yousif et al. ^4, 25^ developed macroscopic Wilson-Cowan^26^ mean-field models on the neuronal network underlying neurological movement disorders, including PD and ET; they implemented the models to approximate the pathological LFP oscillations, including the tremor-range oscillations of ET and the Parkinsonian beta band oscillations. Despite the recognized benefits of these models, they are less effective in replicating firing rate dynamics. Spiking models are more complicated than abstract models, making their parameter fitting mostly based on hand-tuning ^1, 24^. Mean-field models are less effective in reproducing the instantaneous firing rate of a specific nucleus ^3, 4^. In this work, we developed a simple firing rate model to track the *instantaneous* firing rate of stimulated neurons during various frequencies of DBS, and used automatic parameter optimization method to find consistent parameters across different DBS frequencies.

The developed rate model was utilized to mimic instantaneous firing rates of neurons receiving DBS in three basal ganglia and thalamic nuclei, namely, STN, SNr and Vim. We explored the firing rate dynamics in response to different ratios of excitatory and inhibitory presynaptic inputs, during various frequencies of DBS (5 to 200Hz). Similar to Milosevic et al. (2021), the immediate impact of DBS pulses was modeled as inducing synaptic release where synaptic functions were characterized by the Tsodyks & Markram (TM) model ^27^ of short-term synaptic plasticity (STP) ^1^. STP reflects immediately reversible effects of the synapses upon the removal of external stimuli ^1^, and is essentially important in various brain functions, e.g., motor control ^1^, speech recognition ^28^ and working memory ^29^. The integration of STP in the computational modelling can greatly enrich the model’s information processing capability and neuronal behavior predictability ^28^. For the model parameter optimization, we concatenated data across different DBS frequencies, and obtained a single set of optimal model parameters—which is consistent across varying DBS frequencies—for each targeted nucleus. Such consistency in model parameters conformed to the short duration of our DBS recordings (≤ 10s for all DBS frequencies) on each nucleus; the synaptic anatomical structure mostly remains static in response to short (seconds to minutes) external stimuli ^30^. Our developed rate model with the parameter optimization method could accurately reproduce the instantaneous firing rates of various basal ganglia and thalamic neurons receiving different frequencies of DBS. Our model fits well for both synthetic and experimental DBS data. Our work can provide a framework to study the instantaneous effects of DBS parameters on neuronal activity, and may help navigating the DBS parameter space and improve DBS clinical effects.

## MATERIALS AND METHODS

We implemented the same experimental DBS recordings as published in Milosevic et al. (2021)^1^. Thus, the commitment to ethics policies have already been validated ^1^. All experiments conformed to the guidelines set by the Tri-Council Policy on Ethical Conduct for Research Involving Humans and were approved by the University Health Network Research Ethics Board ^1^. Moreover, each patient provided written informed consent prior to taking part in the studies ^1^.

### Method Overview

We developed a rate model to describe the instantaneous firing rate of a population of local neurons receiving DBS. We utilized a sigmoid function to transfer the impact of DBS-induced short-term synaptic plasticity ^1^ to the variability of the firing rate that is expressed by a first order ordinary differential equation (ODE). Specifically, DBS pulses were filtered by the Tsodyks & Markram (TM) model of short-term synaptic plasticity ^27^, and fed to the firing rate differential equation through a sigmoid nonlinear function. In order to estimate the parameters for the nonlinear function and the differential equation, we constructed peristimulus time histograms (PSTH) of recorded spikes (for both synthetic and experimental data) as a reference for modeling instantaneous firing rates. The synthetic data were the simulations from the LIF model in Milosevic et al. ^1^ (see **Supplementary Section 1**), and the experimental data were the single-unit microelectrode recordings (MER) from four basal ganglia and thalamic nuclei—STN, SNr, Vim and reticular thalamic nucleus (Rt)—across specific sets of DBS frequencies in 5∼200Hz. We fitted our rate model to the synthetic data generated by a population of LIF model neurons ^1^. Our simulation study supports model validation by showing its compatibility with an established spiking model (an ensemble of LIF neurons). In both synthetic and experimental data, we obtained the optimal model parameters by minimizing the sum of squared error (SSE) between the model output and the reference PSTH ^31, 32^. For each nucleus, we computed a single set of optimal model parameters that consistently fitted across data from different DBS frequencies {5, 10, 20, 30, 50, 100, and 200Hz}.

### The Computation of Firing Rate with Peristimulus Time Histograms (PSTH)

For each neuron, we simulated or recorded one spike train; simulated (respectively, recorded) spike trains comprised the synthetic data (respectively, experimental data). In either synthetic or experimental data, for each nucleus (STN, SNr, Vim or Rt), we computed the PSTH firing rate (P(t)) by superimposing the spike trains from neurons:

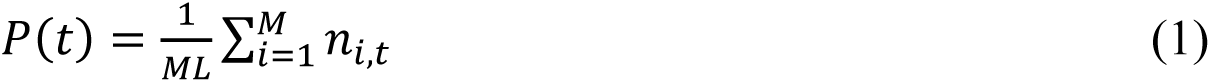

where M is the number of neurons, L (ms) is the length of the PSTH kernel, 𝑛_𝑖,𝑡_ is the number of spikes generated by the i^th^ neuron in the PSTH kernel at time t, i.e., the interval 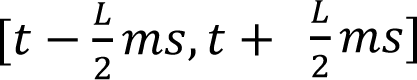. The PSTH firing rate was updated with a time resolution dt = 0.1ms. In terms of the PSTH kernel width, L, we used L = 20ms for both synthetic and experimental data for Vim, Rt, and SNr (consistent with Milosevic et al. (2021) ^1^). For the STN data, we observed high levels of variability (oscillations, see **Results**), thus, a wider PSTH kernel width was needed to effectively represent the firing rate pattern underlying noisy data ^33^. We used L = 50ms for both synthetic and experimental STN data (this kernel width is widely used in computing PSTH ^34, 35^).

### The Synthetic Data

The synthetic data was the firing rate computed with PSTH on 20 spike trains simulated from the LIF model (see **Supplementary Section 1**). We simulated LIF neurons with different DBS frequencies, including {5, 10, 20, 30, 50, 100, and 200Hz}. For each DBS frequency of each nucleus, the simulation time was 1,000ms, with time step of dt = 0.1ms. The simulation stopped at 1 sec because the firing rate steady state was reached before ∼500ms in both synthetic and experimental data, for all nuclei, across different frequencies of DBS (see **Results**). The maximal DBS frequency considered in this work for Vim, Rt, STN, and SNr are 200Hz, 200Hz, 100Hz, and 50Hz, respectively. Experimental data demonstrated that the firing rate of STN and SNr for microelectrode DBS frequencies (using 100µA and symmetric 300µs biphasic pulses) larger than these max frequencies was close to zero ^1^.

### The Experimental Data

The experimental single-unit microelectrode recordings (MER) and data protocols were from Milosevic et al. (2021)^1^. Microelectrodes were used to both deliver DBS and record data, using 100µA and symmetric 300µs biphasic pulses (150µs cathodal followed by 150µs anodal) ^1^. MER has been implemented to optimize the target location for the implantation of DBS electrodes ^36, 37^, as well as the stimulation parameters (frequency, amplitude, pulse width) ^38, 39^. The neuronal firing rate – including both single-unit and mean firing rate – is an important quantifying standard regarding using MER to identify the proper DBS setting and anatomical site ^16, 36, 39, 40^. Following the investigation and localization with MER, macroelectrode is then often implanted to deliver long-time DBS therapy ^37, 41^.

All recordings in STN, SNr and Rt were from patients with Parkinson’s disease, and those for Vim were obtained from patients with essential tremor ^1^. For Vim and Rt, we recorded {5, 10, 20, 30, 50, 100, and 200Hz} DBS data of length {10, 5, 3, 2, 1, 5, and 2s}, respectively. The recording length of the 5Hz DBS data was 5 seconds for STN, and 10 seconds for SNr; we recorded {10, 20, 30, and 50Hz} DBS data of length {5, 3, 2, and 1s} for both STN and SNr. The recording length of the 100Hz DBS data for STN was 3 seconds.

An individual patient received DBS on only one nucleus, whose responses was recorded once for each DBS frequency. For each DBS frequency, we performed 5 to 8 single-unit recordings in different patients (see **Supplementary Table 6** for details) ^1^. In each recording, the narrow stimulus artifacts were removed offline (0.5ms from stimulus onset) ^1^. Then, to detect the spikes, recordings were high pass filtered (≥300Hz) to better isolate the single units, and template matching was done using a principal component analysis method in Spike2 (Cambridge Electronic Design, UK) ^1^. The time stamps of the DBS pulses had small deviations (∼2%) because of the imperfect internal clock of the stimulator ^42^; in the MATLAB script, we adjusted the time stamps so that the DBS pulses were delivered with the accurate frequencies. Using the PSTH formulated in (1), we computed the reference firing rate from these spike trains. The PSTH represents the instantaneous firing rate across spike trains recorded from different patients, and we observed that the data was consistent across patients (see **Results**).

### DBS-induced Input into the Rate Model

The input to our rate model was the DBS-induced post-synaptic current (𝐼_𝑠𝑦𝑛_), and we formulated 𝐼_𝑠𝑦𝑛_ with the Tsodyks & Markram (TM) model on short-term synaptic plasticity (STP) ^27^, as in Milosevic et al. (2021) ^1^. For the neurons receiving DBS, we assumed that each neuron receives inputs from 500 synapses ^1^, and the ratio of the number of excitatory synapses to inhibitory synapses was different for varying nuclei, and shown in **Supplementary Table 1**. The parameters for SNr, Vim, and Rt in **Supplementary Table 1** were the same as those used in Milosevic et al. (2021) ^1^. The excitatory-inhibitory synaptic ratio has high variability in STN neurons ^1, 17^. The synaptic inputs to a minority of STN neurons are dominantly excitatory, whereas the inputs to a majority of STN neurons are dominantly inhibitory ^17^. In this work, we analyzed data recorded from STN neurons receiving evident inhibitory inputs (inhibitory synapses occupies 70%).

Each DBS pulse activated all pre-synaptic inputs simultaneously, and generated DBS-evoked spikes on the presynaptic terminals. The DBS-evoked spikes were filtered by the TM model, generating the post-synaptic current, 𝐼_𝑠𝑦𝑛_, that was obtained by a linear combination of presynaptic excitatory (𝐼_𝑒𝑥𝑐_) and inhibitory (𝐼_𝑖𝑛ℎ_) currents as follows:

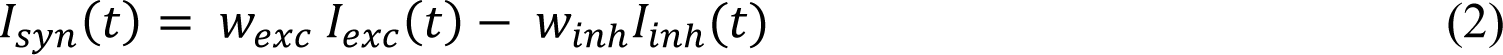

where 𝑤_𝑒𝑥𝑐_ and 𝑤_𝑖𝑛ℎ_ denote the synaptic weights of the modeled excitatory and inhibitory currents, respectively; 𝑤_𝑒𝑥𝑐_ and 𝑤_𝑖𝑛ℎ_ varied for different nuclei ^1^. The values for these weights were summarized in **Supplementary Table 2** (consistent with Milosevic et al. (2021) ^1^, except for STN in which we decreased 𝑤_𝑒𝑥𝑐_to stress the role of inhibitory synapses in fitting STN neurons receiving dominant inhibitory inputs ^17^) .

𝐼_𝑒𝑥𝑐_ (respectively, 𝐼_𝑖𝑛ℎ_) is the total post-synaptic current from all excitatory (respectively, inhibitory) synapses. Each synapse (excitatory or inhibitory) was modeled by the TM model for short-term synaptic plasticity.

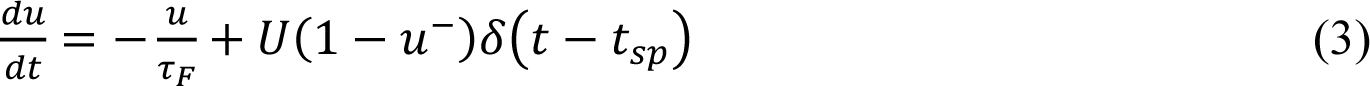

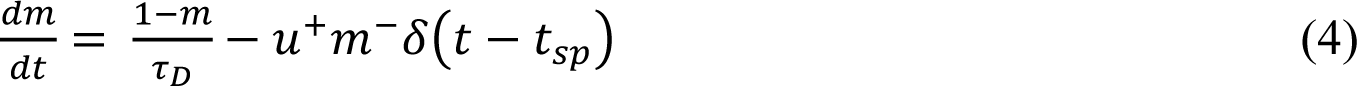

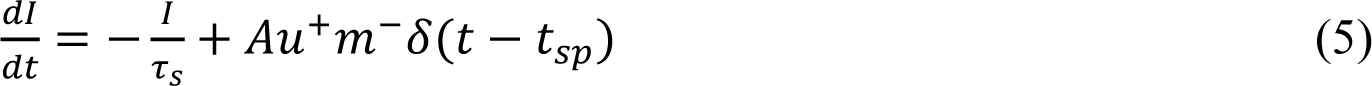

where 𝑢 indicates the utilization probability, i.e., the probability of releasing neurotransmitters in synaptic cleft due to calcium ion flux in the presynaptic terminal. The variable 𝑚 indicates the fraction of available resources after the neurotransmitter depletion caused by neuronal spikes. We denote as 𝑢^−^ and 𝑚^−^ the corresponding variables just before the arrival of the spike; similarly, 𝑢^+^and 𝑚^+^refer to the moment just after the spike. Upon the arrival of each presynaptic spike 𝑡_𝑠𝑝_, 𝑢 increases by 𝑈(1 − 𝑢^−^). If there is no presynaptic activity, 𝑢 exponentially decays to zero; this decay rate is the facilitation time constant, 𝜏_𝐹_. As well, the vesicle depletion process—due to the release of neurotransmitters—was modeled by (4) where 𝑚 denotes the fraction of resources that remains available after the neurotransmitter depletion. In contrast to the increase of 𝑢 upon the arrival of each presynaptic spike, 𝑚 drops and then recovers to its steady state value of 1 (this recovery rate is given by the depression time constant 𝜏_𝐷_). The competition between the depression (𝜏_𝐷_) and facilitation (𝜏_𝐹_) time constants determines the dynamics of the synapse. In the TM model, 𝑈, 𝜏_𝐹_, and 𝜏_𝐷_ are the parameters that determine the types of the synapse, namely, facilitation, pseudo-linear, and depression. The values of the TM model parameters differed across excitatory and inhibitory synapses, and were summarized in **Supplementary Tables 3(A)** & **4(A)**. In (5), 𝐼 and 𝜏_𝑠_ indicate the post-synaptic current and its time constant, respectively. For an excitatory (respectively, inhibitory) synapse, 𝜏_𝑠_ is denoted as 𝜏_𝑒𝑥𝑐_ (respectively, 𝜏_𝑖𝑛ℎ_); these time constants were shown in **Supplementary Table 2**. The absolute response amplitude A = 1 for all situations.

We obtained 𝐼_𝑒𝑥𝑐_(respectively, 𝐼_𝑖𝑛ℎ_) by adding the post-synaptic currents from all excitatory (respectively, inhibitory) synapses. Each nucleus had different proportions of excitatory and inhibitory synapses. Within excitatory (respectively, inhibitory) synapses, the ratio of the 3 types of synapses, namely, facilitation, pseudo-linear, and depression, were also different among different nuclei (**Supplementary Tables 3 and 4**). All TM model parameters in **Supplementary Table 2 – 4** were chosen based on Milosevic et al. (2021) ^1^ and are consistent with the previous modeling works on specific experimental datasets ^43, 44^. Our rate model is robust to the parameters generating the DBS-induced synaptic input (**Supplementary Table 1 – 4**), although the number of these parameters is large (see **Supplementary Section 3**).

### The Rate Model and the Parameter Optimization Method

We used a sigmoid transfer function to link the post-synaptic current (𝐼_𝑠𝑦𝑛_ as defined in (2)) to the rate model on the firing rate of the stimulated nucleus. The rate model underlying a neuronal ensemble receiving DBS was stated as follows:

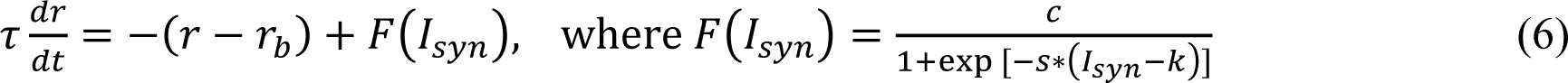

where 𝑟(𝑡) is the neuronal firing rate at time 𝑡, and 𝜏 is the membrane time constant. For the sigmoid transfer function 𝐹(𝐼_𝑠𝑦𝑛_), 𝑐 is the scaling parameter, 𝑠 is the shape parameter, and 𝑘 is the shift parameter. 𝑟_𝑏_ is the baseline firing rate of the modelled nucleus. We confined 𝑟_𝑏_ in biological constraints, based on experimental and synthetic data from both human and mammalian recordings. For Vim, 𝑟_𝑏_ ∈ [10, 50] Hz ^1, 45^; for Rt, 𝑟_𝑏_ ∈ [3, 40] Hz ^1, 22^; for SNr, 𝑟_𝑏_ ∈ [40, 120] Hz ^1, 22^; for STN, 𝑟_𝑏_ ∈ [5, 100] Hz ^1, 46^. The initial value of 𝑟(𝑡) (denoted as 𝑟_𝑖𝑛𝑖_) was computed by simulating a 10s spike train from the LIF model in Milosevic et al. ^1^ with the DBS – OFF condition for the modelled nucleus. 𝑟_𝑖𝑛𝑖_ was computed as: “total number of spikes”/10s; 𝑟_𝑖𝑛𝑖_(Vim) = 39.3Hz, 𝑟_𝑖𝑛𝑖_(Rt) = 5.0Hz, 𝑟_𝑖𝑛𝑖_(SNr) = 57.4Hz, and 𝑟_𝑖𝑛𝑖_(STN) = 27.6Hz. The rate model fit results for Vim, STN and SNr were shown in **Results**; Rt-DBS is a less common choice in clinics to obtain therapeutic effects ^1^, and the corresponding fit result was shown in **Supplementary Figure 2**.

We then inferred the parameter set 𝛷 = {𝜏, 𝑟_𝑏_, 𝑐, 𝑠, 𝑘} by minimizing the sum of squared errors (SSE) between the model output, 𝑟(𝑡), and the reference firing rate, 𝑃(𝑡) (the PSTH firing rate defined in (1)). We fitted the parameters separately for different basal ganglia and thalamus nucleus. For each nucleus, the optimal parameter set 𝛷_𝑜𝑝𝑡_ was the same across all DBS frequencies; such consistency in model parameters conformed to the relatively static synaptic anatomical structure in response to short (seconds to minutes) external stimuli ^30^ (in our case, ≤ 10s for all DBS data).

In the case of both experimental and synthetic data, for the fitting process in the simulation, the sampling resolution was dt = 0.1ms. We ran independent simulations for each DBS frequency and the simulated signal is denoted as 𝑟_𝑓𝑞_(𝛷, 𝑡) = { 𝑟_𝑓𝑞_(𝛷, 𝑡_1_), … , 𝑟_𝑓𝑞_(𝛷, 𝑡_𝑁_)}, which corresponds to a certain DBS frequency (𝑓𝑞) and parameter set 𝛷; *N* is the total number of time points. Similarly, the reference PSTH is denoted as 𝑃_𝑓𝑞_(𝑡) = { 𝑃_𝑓𝑞_(𝑡_1_), … , 𝑃_𝑓𝑞_(𝑡_𝑁_)}. Given the rate model and reference PSTH, the SSE function is:

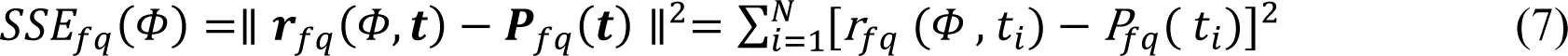

The objective function 𝐽(𝛷) for the parameter optimization was the total SSE across all DBS frequencies in {5, 10, 20, 30, 50, 100, and 200Hz}:

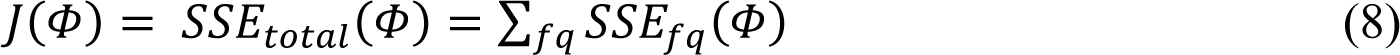

𝐽(𝛷) was minimized with the MATLAB custom function “*fminsearch*”, which used the Nelder – Mead simplex method ^31, 32^ with the 5 variables in the parameter set 𝛷. Starting from an initial point 𝛷_0_ = (𝜏_0_, 𝑟_𝑏,0_, 𝑐_0_, 𝑠_0_, 𝑘_0_), the Nelder – Mead algorithm formed a simplex consisting of 6 vertices around 𝛷_0_. Then the simplex was modified based on 5 operations: reflection, expansion, inside contraction, outside contraction and shrink. In the modified simplex, the algorithm searched for the vertex 𝛷_1_ that minimized the objective function 𝐽(𝛷) and the next iteration started from 𝛷_1_. Compared with the traditional gradient-descent type optimization methods, the advantages of the the simplex method are (i) computation load is reduced because the derivative of the objective function is eliminated; and (ii) the searching direction is not restricted to the local gradient, and the algorithm can quickly approach the minimum in the first few iterations ^32^. We implemented the simplex method in “*fminsearch*” ^32^ to find the optimal parameter set 𝛷_𝑜𝑝𝑡_ = {𝜏_𝑜𝑝𝑡_, 𝑟_𝑏,𝑜𝑝𝑡_, 𝑐_𝑜𝑝𝑡_, 𝑠_𝑜𝑝𝑡_, 𝑘_𝑜𝑝𝑡_} that minimized the objective function 𝐽(𝛷) defined in (8). “*fminsearch*” is an automatic parameter optimization method, which is more accurate and efficient than the commonly used hand-tuning methods used for DBS models ^1, 4, 24^.

## RESULTS

### The Rate Model on Neuronal Dynamics across Multiple DBS Frequencies

We developed a firing rate model that can capture the dynamics of the neuronal activity of varying nuclei across different DBS frequencies. Based on single-unit microelectrode recordings of the neuron receiving DBS in a specific nucleus, we computed the reference firing rate with PSTH. Recordings across DBS frequencies were concatenated, and the optimal parameters were obtained by minimizing the distance to the reference PSTH firing rate (by Equation (8)) across all DBS frequencies using the Nelder – Mead simplex method ^31, 32^. **Figure 1** illustrated our rate model and the parameter optimization method.

**Figure 1.**
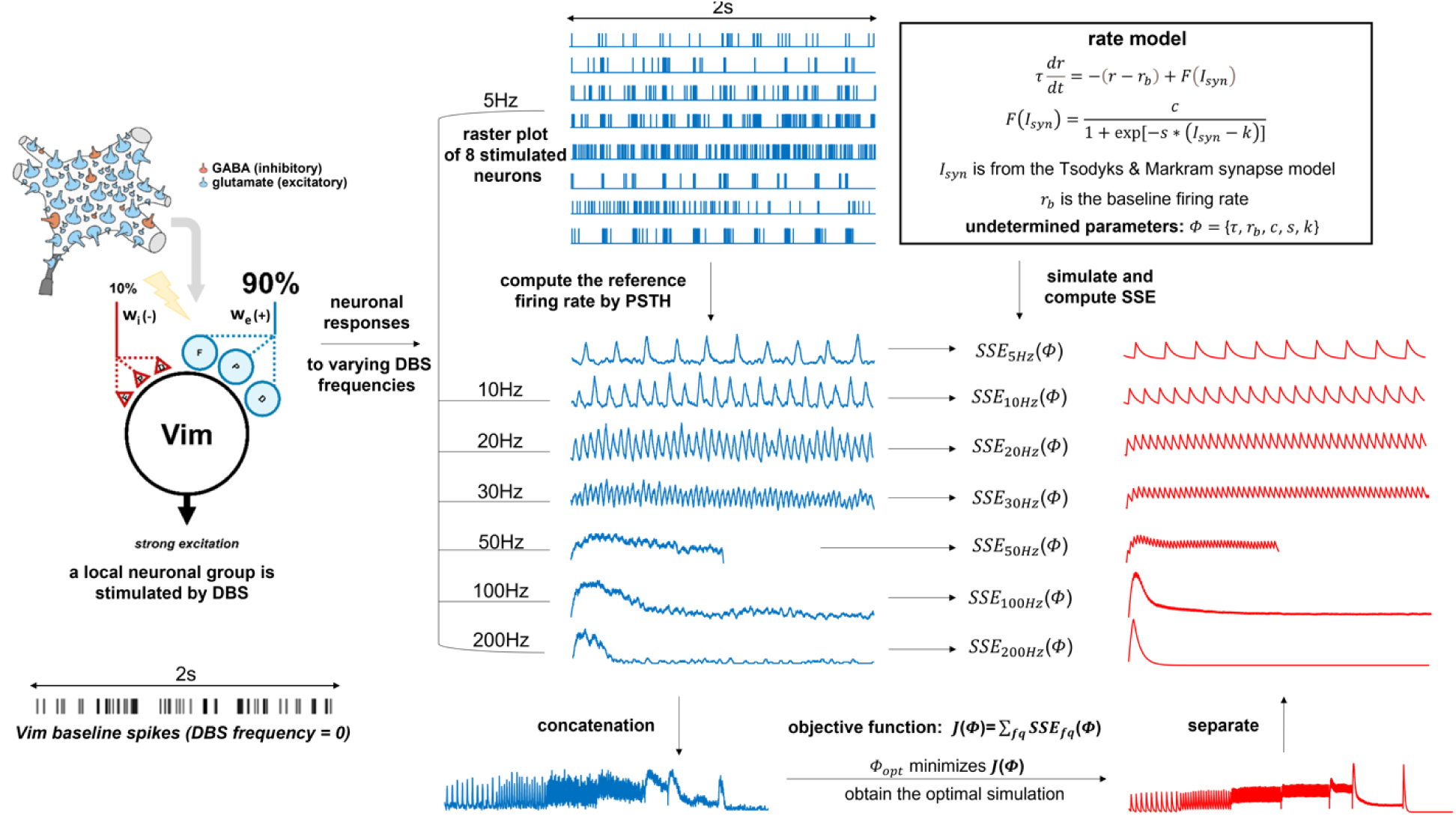
Schematic representation of the rate model (Vim - DBS as the example) A local group of ventral intermediate nucleus (Vim) neurons were stimulated by Deep Brain Stimulation (DBS) ^1^. The 3 synapse types “F”, “P”, and “D” represent “facilitation”, “pseudo-linear”, and “depression”, respectively. For the synapses projected to a Vim neuron, 90% of these synapses release the excitatory (+) glutamate transmitter, and 10% synapses release inhibitory (-) gamma-aminobutyric acid (GABA) transmitter ^1^. We showed the baseline Vim spikes during single-unit microelectrode recording with DBS-OFF. For data from each DBS frequency, we computed the reference firing rate with peristimulus time histogram (PSTH) from the raster plot. Using the rate model, we performed independent simulations for each DBS frequency with the same parameter set 𝛷, and the objective function 𝐽(𝛷) was defined as the total sum of squared error (SSE). We then minimized 𝐽(𝛷) with the Nelder – Mead simplex method, and obtained the optimal parameter set 𝛷_𝑜𝑝𝑡_. We simulated the rate model with 𝛷_𝑜𝑝𝑡_ and obtained the optimal total fit, which minimized the SSE from the concatenated PSTH firing rates. Finally, we separated the total fit and got the optimal fit for each DBS frequency.

### Results — Synthetic Data

We compared firing rates of the rate model simulation with the synthetic data across different DBS frequencies in three different nuclei of the basal ganglia and thalamus, namely, Vim, STN, and SNr. For each nucleus, the synthetic data was the firing rate computed by PSTH with 20 simulated LIF model neurons (see **Materials and Methods**). **Figure 2** showed the model fitted firing rate (red) on top of the reference PSTH firing rate (black) for Vim, STN and SNr across different DBS frequencies. We included a sample spike train of the LIF model neuron to better visualize the relationship between the PSTH and generated spikes.

**Figure 2.**
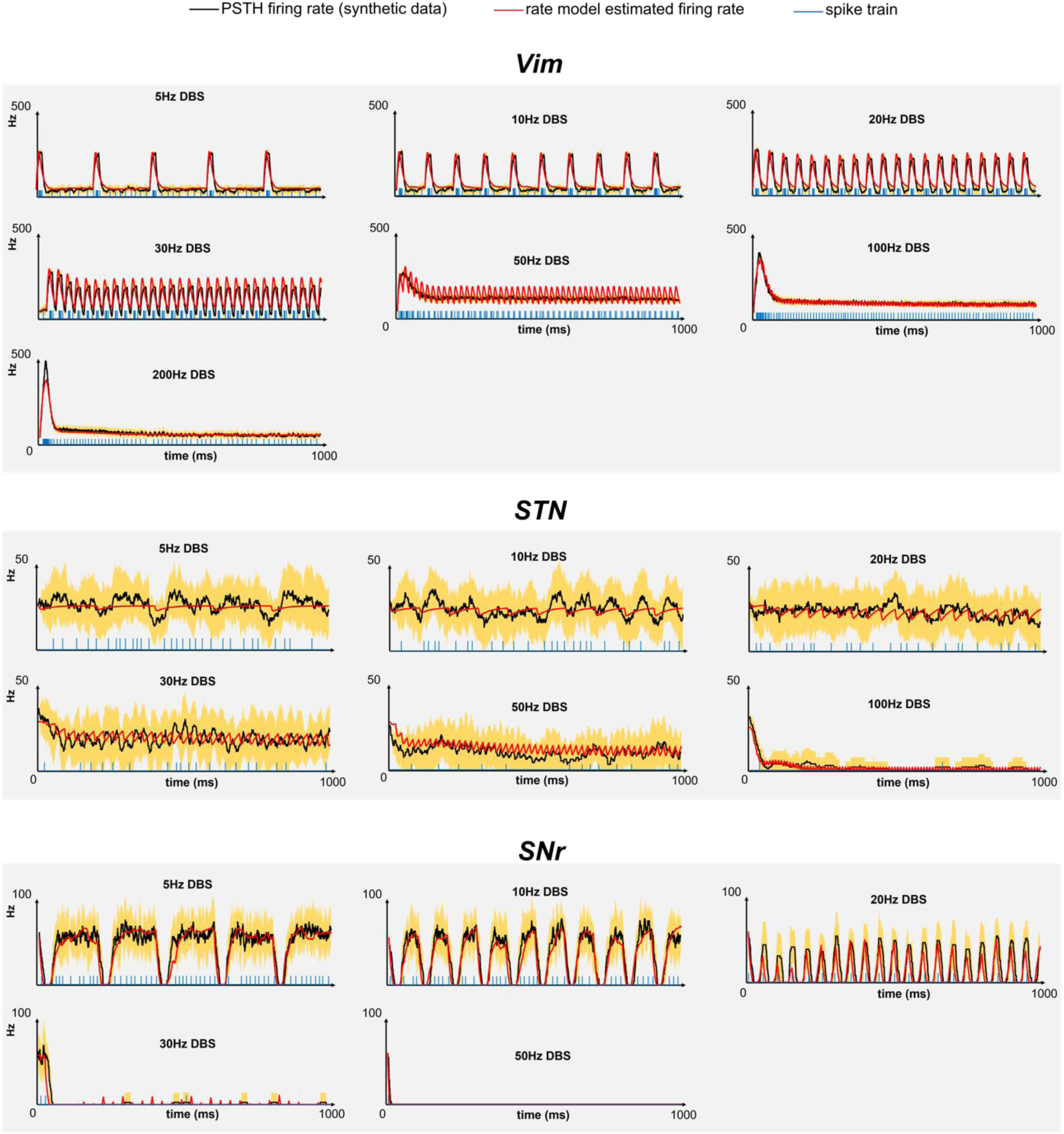
Rate model result for the synthetic data. The rate model was fitted to the synthetic data from the nuclei receiving Deep Brain Stimulation (DBS), in three basal ganglia and thalamic nuclei: ventral intermediate nucleus (Vim), subthalamic nucleus (STN), and substantia nigra pars reticulata (SNr). The synthetic data were the simulated spike trains from the leaky integrate-and-fire (LIF) model in Milosevic et al. ^1^. We compared the firing rate computed by peristimulus time histogram (PSTH) from the synthetic data, firing rate predicted by our rate model and the spike timings from one spike train. The orange shaded region around the reference PSTH was the standard deviation (SD)-based envelope computed from the spike trains; the envelope is Mean ± 1 SD, and is set to be ≥0. DBS stimuli with varying pulse frequencies (5∼200Hz) were delivered to the related basal ganglia and thalamic nuclei. The data from stimulated nuclei receiving lower frequency DBS (5∼50Hz) were recorded in all three nuclei, whereas higher frequency DBS (≥100Hz) was delivered to Vim and STN.

As can be seen in **Figure 2**, the estimated instantaneous firing rate reliably matched the PSTHs generated by ensemble of spiking neurons for all four different nuclei and across all DBS frequencies. In addition, both the steady-state and transient parts of the neurons’ firing rate were replicated using the rate model. For each neuron, given each DBS frequency, we plotted the standard deviation (SD) envelope (**Figure 2**, orange shaded region) – based on the simulated spike trains – around the reference (total) PSTH firing rate. For each spike train, we computed the corresponding instantaneous firing rate with the same kernel as that used for calculating the reference PSTH. The SD envelope of the STN data was wide, demonstrating higher level of variability compared to other nuclei (**Figure 3**). We found that model predicted firing rate lied almost always within the corresponding SD envelopes for all neurons (**Figure 2**), confirming the reliability of the model for the synthetic data.

**Figure 3.**
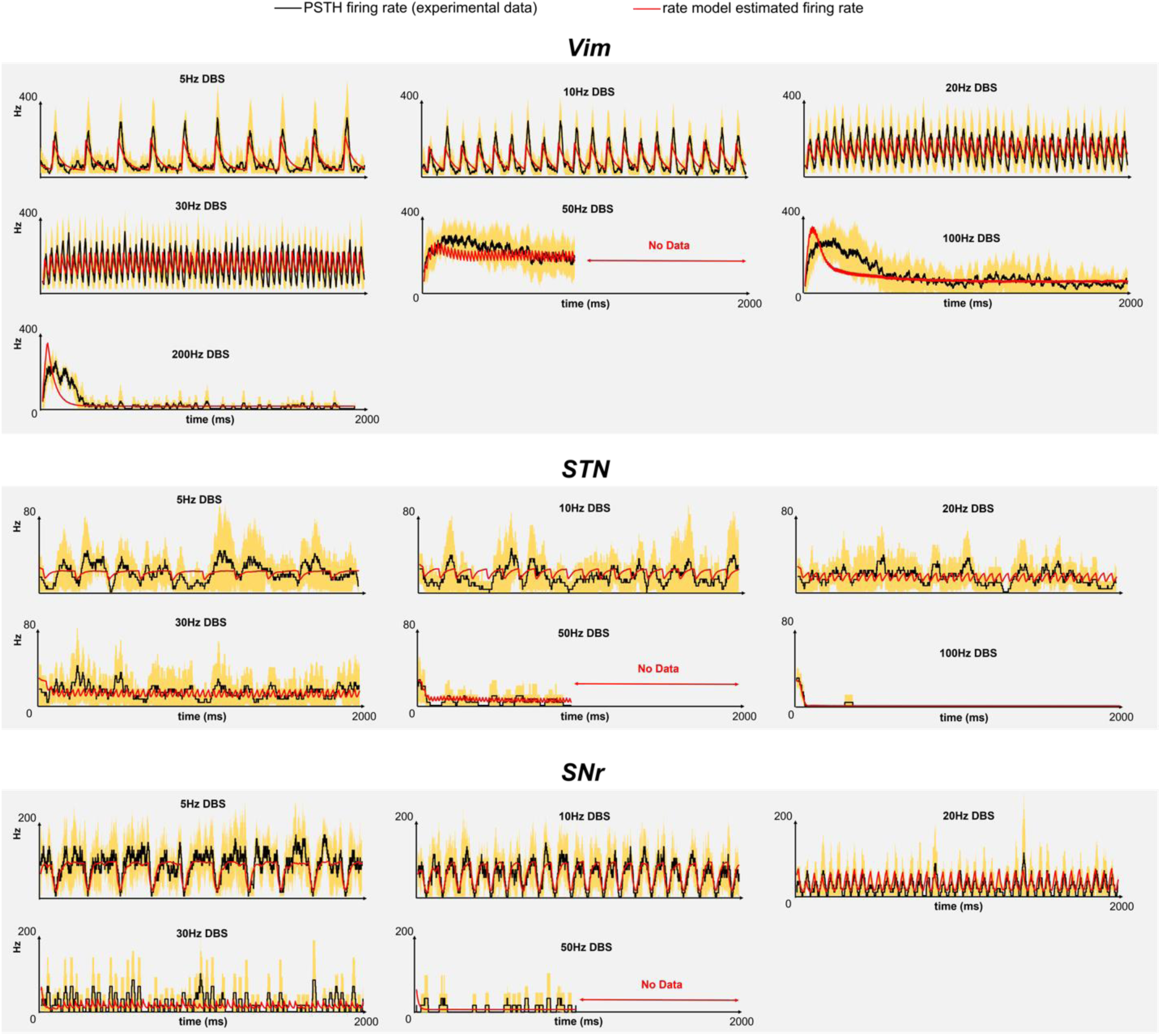
Rate model result for the experimental data. The rate model was fitted to the experimental data (single-unit microelectrode recordings) from the nuclei receiving Deep Brain Stimulation (DBS), in three basal ganglia and thalamic nuclei: ventral intermediate nucleus (Vim), subthalamic nucleus (STN), and substantia nigra pars reticulata (SNr). We compared the firing rate computed by peristimulus time histogram (PSTH) from the experimental data and the firing rate predicted by our rate model. The experimental data were from single-unit microelectrode DBS recordings in each basal ganglia and thalamic nucleus. The recording length for 50 Hz DBS data was ∼1s. The data from lower frequency DBS (5∼50Hz) were recorded in all 3 nuclei, whereas higher frequency DBS (≥100Hz) was delivered to Vim and STN. The orange shaded region around the reference PSTH shows the standard deviation (SD)-based envelope computed with the spike trains from the experimental recordings; the envelope is Mean ± 1 SD, and is set to be ≥0.

We fitted the rate model parameters based on the synthetic data from DBS frequencies {5, 10, 20, 30, 50, 100, and 200Hz}. In order to fully validate the rate model, and justify its generalizability to unobserved DBS frequencies, we used the existing rate model parameters to predict the synthetic data from a broader choice of DBS frequencies. The mean and sample standard deviation of the normalized mean squared error (NMSE) of the rate model fit were presented in **Table 1**.

**Table 1:** Normalized mean squared error (NMSE) of rate model fit to synthetic data from different DBS frequencies

|  | Vim |  | STN |  | SNr |  |
| --- | --- | --- | --- | --- | --- | --- |
|  | observed frequencies | all frequencies | observed frequencies | all frequencies | observed frequencies | all frequencies |
| mean | 7.7% | 4.6% | 8.5% | 11.8% | 11.1% | 9.8% |
| standard deviation | 4.8% | 4.5% | 7.9% | 11.9% | 11.8% | 9.5% |

In **Table 1**, the “observed frequencies” represented the DBS frequencies {5, 10, 20, 30, 50, 100, and 200Hz} used for fitting the rate model parameters, which were implemented to predict the synthetic data from “all frequencies”, including {2.5, 5, 7.5, 10, 15, 20, 30, 40, 50, 60, 70, 80, 90, 100, 110, 120, 130, 140, 150, 160, 170, 180, 190, and 200Hz} (for STN and SNr, the maximum is 100Hz and 50Hz, respectively). The NMSE from all DBS frequencies was small in Vim, STN and SNr (**Table 1**); this showed that the rate model can accurately predict the synthetic data. The NMSE between observed and all frequencies was consistent (**Table 1**); thus, the rate model parameters fitted to data from observed DBS frequencies can be consistently generalized to data from both observed and unobserved DBS frequencies.

To conclude, these above results indicated that our rate model was compatible with the established LIF model in Milosevic et al. (2021) ^1^. Furthermore, we also demonstrated that a simple abstract model (our rate model) can efficiently replicate the results from simulating an ensemble of LIF neurons.

### Results — Experimental Data

To test the potential of the rate model in reproducing instantaneous firing rate of an ensemble of neurons recorded from human brain during DBS, we fitted the rate model to experimental data obtained from single-unit microelectrode recordings ^1^ on three different nuclei of basal ganglia and thalamus: Vim, STN, and SNr. To calculate the firing rate of an ensemble of neurons for each nucleus, spikes recorded from 5 to 8 different individuals were combined, and the instantaneous firing rate was then obtained by calculating the PSTH. Similar to the results for the synthetic data, we estimated model parameters that fitted the rate model output to the PSTH. **Figure 3** showed the results of the fit model output with the PSTH for different nuclei and different DBS frequencies.

Firing rates estimated by the rate model track both transient-(mostly apparent for DBS frequencies ≥30 Hz) and steady-state components of PSTH’s of all nuclei. The NMSE computed based on the concatenated signal from all DBS frequencies for Vim, STN and SNr is 14.1%, 18.1% and 9.5%, respectively. Similar to synthetic data (**Figure 2**), we plotted the SD envelope (**Figure 3**, orange shaded region) of experimentally recorded spike trains – around the reference (total) PSTH firing rate. As can be observed in this figure, the model predicted firing rate can track neuronal responses of different substructures of the basal ganglia and thalamus in response to different DBS frequencies (model output lies, almost always, within the corresponding SD envelopes (**Figure 3**)).

However, the rate model could not reproduce the transient response of the Vim for DBS frequencies of 100 Hz and 200 Hz as accurately as it could for other nuclei. The rate model generated a shorter transient response compared to that observed in experimental recordings. This mismatch between transient responses might be originated from neuronal network effects and was further addressed in **Discussions**. Besides, high variabilities observed in the STN data were reflected by the wide SD envelope (**Figure 3**). It is to be noted that the large variabilities in STN data were not accurately captured by the model and such variabilities might be induced by the network effects from external globus pallidus (GPe) ^47, 48^ (see **Discussions**).

### Physiological Implications of the Model Parameters

The optimal rate model parameters, 𝛷_𝑜𝑝𝑡_ = {𝜏_𝑜𝑝𝑡_, 𝑟_𝑏,𝑜𝑝𝑡_, 𝑐_𝑜𝑝𝑡_, 𝑠_𝑜𝑝𝑡_, 𝑘_𝑜𝑝𝑡_}, of each nucleus, for both synthetic and experimental data, were listed in **Table 2**:

**Table 2:** Rate model optimal parameters (3 significant figures)

| source | site | $\tau_{opt}$ | $r_{b,opt}$ | $c_{opt}$ | $s_{opt}$ | $k_{opt}$ |
| --- | --- | --- | --- | --- | --- | --- |
| synthetic data | Vim | 10.4 | 10.0 | 433 | $4.40 \times 10^{-3}$ | 616 |
|  | STN | 36.0 | 27.5 | -51.5 | -0.470 | -14.0 |
|  | SNr | 11.1 | 77.1 | -96.6 | -0.273 | -17.8 |
| | Rt | 11.9 | 2.53 | 392 | $3.20 \times 10^{-2}$ | 112 |
| experimental data | Vim | 45.1 | 13.4 | 687 | $5.81 \times 10^{-2}$ | 548 |
|  | STN | 24.7 | 27.6 | -34.4 | -0.425 | -5.42 |
|  | SNr | 11.5 | 95.3 | -88.6 | -0.236 | -21.5 |
|  | Rt | 32.2 | 3.00 | 578 | 32.9 | 53.3 |

The optimal model parameters were in distinct ranges for different nuclei of the basal ganglia and thalamus. For the same nucleus, the optimal parameters of synthetic and experimental data were generally similar but differences existed. In fitting the model to experimental data, we observed that model parameters were robust to the data variability caused by removing data of an individual patient (i.e., leave-one-out fits, see **Supplementary Section 4**). The time constant (𝜏_𝑜𝑝𝑡_) and the steady-state value (i.e., baseline firing rate 𝑟_𝑏,𝑜𝑝𝑡_) represent the dynamics of the underlying system. The DBS impact on the firing rate dynamics was characterized through the non-linearity 𝐹(𝐼_𝑠𝑦𝑛_) with the parameters {𝑐_𝑜𝑝𝑡_, 𝑠_𝑜𝑝𝑡_, 𝑘_𝑜𝑝𝑡_}. The combination of the parameters in **Table 2** forms the firing rate response to the DBS-induced input synaptic current 𝐼_𝑠𝑦𝑛_ through the sigmoid function as defined in (6): 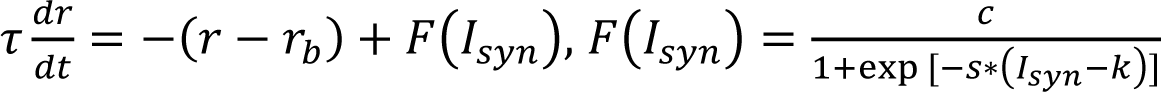 To better assess the effect of these parameters on the rate model, we plotted the model estimated firing rate (𝑟) versus 𝐼_𝑠𝑦𝑛_in **Figure 4** to detect the firing rate dynamics of each nucleus across varying DBS frequencies. In other words, this figure described the *f*-*I* curve for each nucleus receiving DBS. The *f*-*I* curve was created by the estimated parameters in **Table 2** for both synthetic and experimental data. The firing rate and 𝐼_𝑠𝑦𝑛_were ordered according to their percentiles (from 0 to 100), and the percentiles were matched at each point in the curve to qualitatively represent the relationship between the firing rate and input current ^49, 50^. A similar curve was created in Lim et al. (2015) ^20^ for the rate network model to calculate the current-rate transfer function. In our proposed model, see (6), the sigmoid transfer function 𝐹(𝐼_𝑠𝑦𝑛_) modeled the firing rate variability caused by the input synaptic current (𝐼_𝑠𝑦𝑛_); see **Supplementary Figure 1** for 𝐹(𝐼_𝑠𝑦𝑛_) in each nucleus across different DBS frequencies.

**Figure 4.**
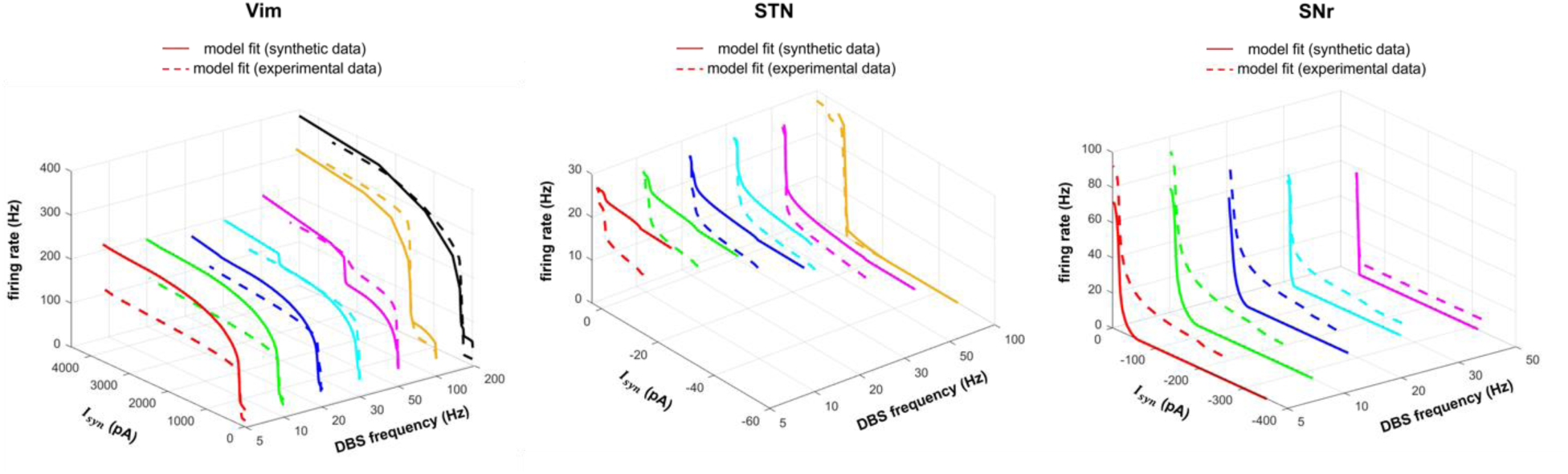
Rate model estimated firing rate in response to the input synaptic current. The rate model was fitted to both synthetic and experimental data; the synthetic data were from the spiking model in Milosevic et al. (2021) ^1^, and the experimental data were single-unit microelectrode recordings. We presented the model fits across all Deep Brain Stimulation (DBS) frequencies, in each of the three basal ganglia and thalamic nuclei: ventral intermediate nucleus (Vim), subthalamic nucleus (STN), and substantia nigra pars reticulata (SNr). We plotted the relationship between the rate model estimated firing rate and the simulated input synaptic current (𝐼_𝑠𝑦𝑛_) to the nuclei receiving DBS. The unit of 𝐼_𝑠𝑦𝑛_ is picoampere (pA) ^51, 52^. Each point in a curve represents that the corresponding firing rate and the input synaptic current have the same percentile (in the range 0 to 100).

In **Figure 4**, the model estimated firing rate monotonically increases as the input synaptic current becomes more excitatory or less inhibitory, in both synthetic and experimental data. Note that the minus sign of 𝐼_𝑠𝑦𝑛_indicates inhibition and the positive sign denotes excitation. For SNr and STN with predominantly inhibitory inputs, the *f-I* curve is monotonically decreasing, across all DBS frequencies, indicating that neurons in this nucleus received more inhibitory synaptic inputs when DBS is ON. When DBS is active, the firing rate quickly drops from its baseline value given negative net current received by SNr neurons. The slope of this fast decay is facilitated in higher frequencies of DBS. Unlike SNr neurons that receive more inhibitory synaptic inputs given DBS pulses, the firing rate of Vim neurons increases monotonically with respect to total synaptic current whose net effect is positive when DBS is ON. The slope of the *f-I* curve increases for higher frequencies of DBS in Vim neurons, and the model simulated firing rate saturates when 𝐼_𝑠𝑦𝑛_is large. One can note that in experimental and synthetic data, for each nucleus, the calculated *f-I* curves (**Figure 4**) were generally similar, although some differences existed. For the data from a certain DBS frequency in one nucleus, the mismatch between the *f-I* curves represents the difference between synthetic and experimental data (**Figure 4**). In future works, we will explore the reason of such mismatch, and work on improving the model generating the synthetic data to obtain more accurate simulations.

## DISCUSSIONS

In this paper, we developed a firing rate model to replicate instantaneous firing rate of three nuclei in the basal ganglia and thalamus, namely, Vim, STN and SNr, during DBS. The rate model enabled accurate estimation of the firing rate dynamics obtained from both synthetic and experimental data. Compared with the spiking network models ^24, 53^, the rate model has fewer number of equations and lower dimensionality in model parameters, thus, making the rate model much more efficient for automatic parameter estimation ^54^. For replicating the firing rate of each stimulated nucleus, we optimized a single set of model parameters fitted consistently across various DBS frequencies, including both observed frequencies (for model training) and unobserved frequencies.

### Methods for Modeling the Impact of DBS

In this work, we modeled the DBS impact as inducing synaptic release, in which the synaptic function was characterized by the Tsodyks & Markram (TM) model ^27^ for short-term synaptic plasticity (STP) ^1^. Such modeling strategy on the DBS effect is consistent with Farokhniaee and McIntyre (2019) ^14^ as well as our previous work Milosevic et al. (2021) ^1^, where the TM model was utilized to produce DBS-induced post-synaptic current.

It is to be noted that there are other methods to model the impact of DBS. For example, Schmidt et al. (2020) modeled the DBS effect as generating a spherical electrical field that affects the potential of all neuronal elements (including soma, axon, etc.) within a certain distance from the DBS electrode ^55^. Farokhniaee and Lowery (2021) modeled the effect of the subthalamic-DBS on cortical neurons through the hyperdirect pathway by inducing changes in the post-synaptic membrane potential through a delta-function ^24^. Farokhniaee and Lowery (2021) did not model the DBS effect as inducing synaptic release as the electrical field generated by subthalamic-DBS did not directly activate synapses that were projected onto cortical neurons ^24^. The electrical field induced by DBS can be non-spherical if using multiple electrodes or directional leads ^56, 57^. In future works, we will compare the impacts (electrical fields) induced by different DBS electrodes, and formulate these impacts as inducing synaptic release characterized by the TM model.

### Further Discussions on Model Physiological Implications

For Vim neurons, the large excitatory 𝐼_𝑠𝑦𝑛_ (**Figure 4**) was originated by the high proportion of glutamatergic pre-synaptic inputs ^1^; the firing rate generally increases as we stimulate with higher DBS frequency. The modeled predicted firing rate from either experimental or synthetic data (**Figure 4**) has similar patterns across different DBS frequencies, based on the amplitude of 𝐼_𝑠𝑦𝑛_: (i) When 𝐼_𝑠𝑦𝑛_ is small, the firing rate increases quickly as the excitatory input increases; (ii) When 𝐼_𝑠𝑦𝑛_ is large, the firing rate slowly increases and finally saturates when 𝐼_𝑠𝑦𝑛_ is large.

For SNr (**Figure 4**), we observed that the firing rate is low and close to zero in most of the range of 𝐼_𝑠𝑦𝑛_, in both experimental and synthetic data across all DBS frequencies. The modeled firing rate (**Figure 4**) has similar patterns across different DBS frequencies, based on the 𝐼_𝑠𝑦𝑛_amplitude: (i) when the inhibition is strong (i.e., large 𝐼_𝑠𝑦𝑛_amplitude), the firing rate is low and mostly close to 0; (ii) when the inhibition is removed (i.e., 𝐼_𝑠𝑦𝑛_∼ 0), the firing rate returns to the baseline at ∼60Hz ^1^. The large inhibitory 𝐼_𝑠𝑦𝑛_is consistent with the high proportion (∼90%) of gamma-aminobutyric acid (GABA) pre-synaptic inputs to SNr neurons ^1^.

In **Supplementary Figure 1**, we showed that the sigmoid nonlinearity 𝐹(𝐼_𝑠𝑦𝑛_) in response to 𝐼_𝑠𝑦𝑛_ is generally similar for synthetic and experimental data, for all three substructures (Vim, STN and SNr). For the data from a certain DBS frequency in one nucleus, the mismatch between F(I_syn_) represents the difference between synthetic and experimental data (**Supplementary Figure 1**). For SNr and STN (dominant inhibitory), 𝐹(𝐼_𝑠𝑦𝑛_) is mostly negative with the evidently inhibitory 𝐼_𝑠𝑦𝑛_. For Vim, as 𝐼_𝑠𝑦𝑛_ increases, 𝐹(𝐼_𝑠𝑦𝑛_) first increases then plateaus; this indicates that the increase of the firing rate is restricted (see the saturation in **Figure 4** **– Vim**), although the postsynaptic current (𝐼_𝑠𝑦𝑛_) is large. Since 𝐼_𝑠𝑦𝑛_ incorporates the STP rules in the synapse, 𝐹(𝐼_𝑠𝑦𝑛_) partially reflects the firing rate regulation of the neuron beyond the synaptic contacts.

### Limitations and Future Work

Our model represented local groups of neurons, but we did not model the network interactions among the connected neuronal groups. For example, the afferent synaptic connections to Vim neurons consist of both excitatory and inhibitory effects ^4^. For a Vim neuron, the excitatory synapses are mainly from the cortex and the cerebellum ^1^, and the inhibitory synapses are mainly from Rt and the interneurons ^58, 59^. We modelled the local Vim neurons receiving DBS, whereas the DBS-induced upstream / downstream network effects from other nuclei were not included in the model.

From **Figure 3** (experimental data, Vim), we saw that the firing rate prediction with our model is not very accurate at high DBS frequencies (100Hz and 200Hz). In Vim experimental data at high frequency DBS, the initial transient large firing rate response (∼400ms) was clearly longer than the model simulation (∼200ms). We hypothesize that the model firing rate predictions in response to DBS at high frequencies can be better explained if we include network effects into the rate model in this work.

Results for STN-DBS (experimental data) in **Figure 3** showed that our model cannot capture oscillations in the firing rate, in particular for the 5Hz and 10Hz DBS frequency. Those oscillations in STN are mainly caused by interaction of inputs from the cortex ^60^ and the external globus pallidus (GPe) ^47^. In particular, the STN-GPe recurrent network contributes to the oscillatory behaviors observed in STN ^47, 48^. Thus, in order to completely model the firing rate oscillations in STN, a model with a single equation, representing dynamics of a single population of stimulated neurons, may not be sufficient. We anticipate that the fitting accuracy to STN oscillations will be improved by incorporating the related network effects.

Synaptic inputs to STN neurons are diverse; some STN neurons receive predominantly excitatory whereas other STN neurons receive predominately inhibitory inputs ^1, 15, 17^. In this work, we only modelled STN nucleus receiving dominant inhibitory inputs, that have been shown to be persistent at high frequency DBS in both rat ^15^ and human ^17^. In the future work, we aim to incorporate STN neurons that receive predominantly excitatory synapses, that have been shown to be governed by different short-term synaptic depression (STD) dynamics compared to inhibitory inputs to STN ^15^. Furthermore, incorporating both dynamics of inhibitory and excitatory inputs to STN will help us predict the firing rate of STN more precisely.

The DBS experimental data in this work was recorded in a relatively short time scale (≤10s) for each stimulation frequency (5∼200Hz). In response to external stimuli with a longer time scale (≥ minutes), the stimulated neurons may exhibit features associated to long term synaptic plasticity (LTP), which is the long-term change in the synaptic connectivity or morphology ^61–63^. The LTP is widely observed in cortical neuronal circuits on memory ^62^, learning ^64^ and neuromodulations ^64, 65^, etc. The modeling on the possible LTP during long-time DBS will be addressed in our future works.

In our model, the instantaneous firing rate, for each substructure, was calculated from spikes recorded from different individuals. Using the standard deviation envelope in **Figure 3**, we observed that the data from different individuals were generally consistent although some variabilities (among individuals) also exist. One should notice that although our modeling framework is generalizable for stimulated neurons, specific features (e.g., variability) of single neurons are not fully represented in the abstract model. This is a limitation of the current study. In future studies, we aim to conduct multiple recordings, for each frequency of DBS, from each participant (i.e., patients with PD and tremor) and explore underlying physiological features in our model.

In clinical applications, an appropriate choice of DBS parameters, in particular for DBS frequencies, is critical for successful therapeutic results and reducing side effects ^12, 66^. However, the process of choosing DBS parameters relies largely on inefficient trial-and-error approaches that are based on neurologist experience and clinical observations ^4, 12, 67^. Based on the relationship between neuronal pathological firing rates and the disease symptoms, our rate model has a potential to provide a quantitative method for choosing the optimal DBS frequency that is therapeutically effective (i.e., symptom relieving). The determination of the optimal DBS parameters depends on the further understanding of the neuronal network mechanisms underlying the disease and the therapeutic effects of DBS ^4^. Using results of the present study, we, in our future works, will develop firing rate network models to detect the mechanisms underlying the interacting neuronal circuits influenced by DBS. Neuronal network models can be implemented in closed-loop control systems to automatically adjust DBS frequencies based on the needs of individual patients ^66, 68^.

It is noted that we focused on tuning DBS frequencies in this work; although the DBS frequency is the most commonly tuned parameter in clinical applications ^69^, the optimization of other DBS parameters (e.g., pulse width and amplitude) may also contribute to a better clinical performance. Even with the limitations in not modeling DBS pulse width and amplitude, our model fitted the clinically recorded experimental data consistently across varying DBS frequencies, for each nucleus – STN, SNr, Vim and Rt.

## Conclusion

In this paper, we developed a firing rate model on the basal ganglia and thalamic neurons receiving DBS. We optimized the consistent model parameters that can be effectively fitted across different stimulation frequencies of DBS. The optimal rate model parameters characterize the DBS impact on the firing rate dynamics, and can quantitatively predict the firing rate patterns induced by both low and high frequency DBS. Our model could accurately reproduce the firing rate obtained from both synthetic data and experimental single-unit microelectrode recordings (MER). Such consistency and accuracy of the model fits demonstrated that our model can infer the DBS effects on neuronal activity, and potentially be further implemented in optimizing the DBS frequency to improve the clinical performances of DBS.

### Data Availability Statement

The human experimental datasets were from Milosevic et al. (2021) ^1^. The codes to regenerate the results can be accessed online from https://github.com/nsbspl/DBS_Mechanism_Cellular.

## SUPPLEMENTARY INFORMATION

### Supplementary Section 1 — The Leaky Integrate-and-Fire (LIF) Spiking Model

The LIF model in Milosevic et al. (2021) ^1^ was implemented to generate the synthetic data used in our analysis. In this LIF model, the total input current (𝐼_𝑡𝑜𝑡𝑎𝑙_) to a stimulated neuron consisted of two components: the post-synaptic current (𝐼_𝑠𝑦𝑛_) and background noise (𝐼_𝑛𝑜𝑖𝑠𝑒_), i.e.

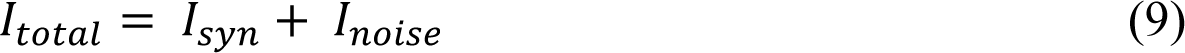

### 𝐼_𝑠𝑦𝑛_ was obtained with the Tsodyks & Markram model formulated in **Materials and Methods**

𝐼_𝑛𝑜𝑖𝑠𝑒_ was modeled by the Ornstein-Uhlenbeck (OU) process with time constant of 5ms ^70^, and is written as:

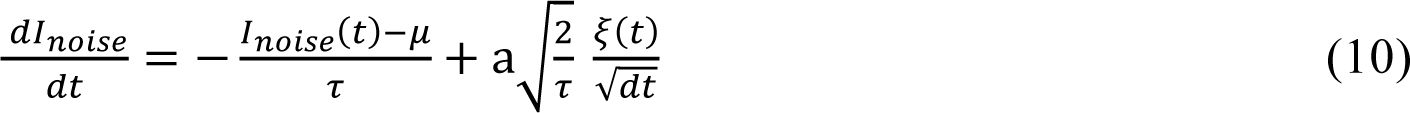

where 𝜉(𝑡) is white noise with mean = 0 and variance = 1. 𝜏 = 5ms is the time constant, 𝜇 and a indicate the mean and standard deviation of 𝐼_𝑛𝑜𝑖𝑠𝑒_, respectively. {𝜇, a} was different for each nucleus; the different values were shown in **Supplementary Table 5**.

The dynamics of the membrane potential of a stimulated neuron in an LIF model can be written as:

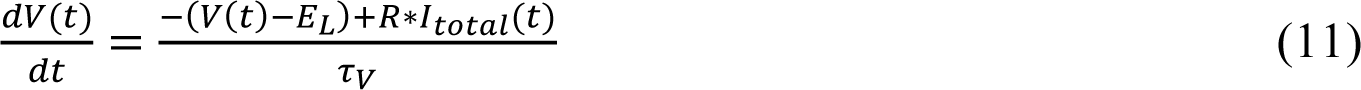

where 𝐸_𝐿_= −70mV is the equilibrium potential, R (scaling parameter) = 1, and 𝜏_𝑉_ (membrane time constant) = 10ms. 𝐼_𝑡𝑜𝑡𝑎𝑙_ is the total input current (Equations (9) – (10)). Spikes occur when 𝑉 ≥ 𝑉_𝑡ℎ_, for 𝑉_𝑡ℎ_ = −40mV. The reset voltage is −90mV and the absolute refractory period is 1ms.

### Supplementary Section 2 — Comparison with a Common Simplified Approach to Optimize DBS Model Parameters

The conventional approach for modeling DBS-evoked neural response—regardless of the optimization technique or the modeling framework—was based on data recorded from a single DBS frequency ^4, 25^, less stressing the frequency-dependent behavior of DBS neural responses. High frequency DBS was considered as clinically effective and used to fit the model ^4^; however, therapeutic effective DBS frequencies are often undetermined and depend on individual situations ^1, 71, 72^. In some certain case studies, the effective DBS frequencies were ∼130Hz for Vim ^73^, ≥ 100Hz for STN ^69^ and below 70Hz for SNr ^74^. Unlike other studies, we developed a “concatenated-frequencies method” in optimizing model parameters, by incorporating the contribution of various DBS frequencies in our rate model and estimated model parameters that are consistent across those frequencies. The concatenated-frequencies method included all the previously mentioned DBS frequencies, i.e., {5, 10, 20, 30, 50, 100, and 200Hz} (for STN and SNr, the maximum is 100Hz and 50Hz, respectively). To compare the concatenated-frequencies method with the parameter estimation based on only a single DBS frequency (“single-frequency method”), we fitted our model using a single DBS frequency for Vim-DBS = 100Hz, STN-DBS = 100Hz and SNr-DBS = 20Hz; the comparison result was shown in **Supplementary Figure 3**. For both optimization methods, we used the estimated model parameters to replicate the firing rate for 24 observed and unobserved DBS frequencies in 0∼200 Hz, including {2.5, 5, 7.5, 10, 15, 20, 30, 40, 50, 60, 70, 80, 90, 100, 110, 120, 130, 140, 150, 160, 170, 180, 190, and 200Hz} (for STN and SNr, the maximum is 100Hz and 50Hz, respectively). We used the normalized mean squared error (NMSE) to measure the error between the estimated firing rate and the reference firing rate computed by PSTH. **Supplementary Figure 3(A)** showed a sample of estimated firing rates using single-frequency and concatenated-frequencies methods for Vim. For the parameter estimation based on a single DBS frequency, model parameters were obtained from Vim-DBS = 100Hz and the estimated firing rate was plotted for DBS of 5Hz & 100Hz. The estimated firing rate based on a single DBS frequency worked well for that frequency but failed for the other (5Hz). However, the estimated firing rate based on multiple DBS frequencies reliably reproduced both the transient- and steady-states of the instantaneous firing rate. For the parameter estimation based on a single DBS frequency, we observed (data not shown) that the estimated firing rate could only replicate the original instantaneous firing rates for DBS frequencies of 100Hz and 50Hz; it produced large deviations for smaller DBS frequencies.

The NMSE calculated using single and multiple DBS frequencies were shown in **Supplementary Figure 3(B)** for all nuclei. The NMSE’s calculated by multiple DBS frequencies were significantly smaller (with regard to mean and standard deviation) than those obtained by a single DBS frequency (**Supplementary Figure 3(B)**). The small NMSE compared to that based on single DBS frequency indicated that our proposed rate model with the concatenated-frequencies optimization method could much better reproduce the PSTH firing rate (ANOVA, p < 0.05 for all nuclei; p = 8.45 × 10^-3^ for Vim, p = 4.66 × 10^-4^ for STN, p = 0.0286 for SNr).

### Supplementary Section 3 – Input Parameter Sensitivity Analysis

The sensitivity analysis was performed consistent with Farokhniaee and McIntyre (2019) ^14^ for all synaptic input parameters shown in **Supplementary Tables 1 – 4**. Each parameter was chosen randomly within a ±10% error range; specifically, for a given parameter value *m*, a new parameter was chosen randomly in [0.9m, 1.1m]. In **Supplementary Tables 1**, we randomized the proportion of inhibitory synapses, and the remaining was the proportion of excitatory synapses. In **Supplementary Tables 3(B) and 4(B)**, we randomized the proportion of facilitation and pseudo linear synapses, and the remaining was the proportion of depression synapses. All parameters in **Supplementary Tables 2, 3(A) and 4(A)** were randomized. In total, we randomized 27 parameters in the sensitivity analysis.

As a proof of concept, we applied the sensitivity (for parameters underlying the input signal) to Vim neurons. We randomized 10 sets of synaptic input parameters. For each set of parameters, we simulated the rate model in response to different DBS frequencies, ranging from 5 Hz to 200 Hz (frequency step = 5 Hz), using the settings {τ, r_b_, c, s, 𝑘} in **Table 2** that characterized the Vim experimental data. Each simulation lasted 1s, and we computed the mean firing rate during the simulation. The results were shown in **Supplementary Figure 4**, in which the red trace indicates the synaptic input parameters used in this study. In **Supplementary Figure 4**, we can see that the traces with randomized synaptic input parameters are majorly within ±20% error range of the mean firing rate, showing that the rate model is reasonably robust to the change of synaptic input parameters (randomizing a large number of synaptic input parameters together).

### Supplementary Section 4 – Leave-one-out Validation in Fitting Experimental Data

We performed the leave-one-out validation to test the robustness of the rate model fit to the variability caused by leaving spikes recorded from an individual neuron. Note that the variability in experimental recordings from individual neurons was demonstrated as the standard deviation (SD)-based envelope shown in **Figure 3**. As a proof of concept, we performed the model validation on Vim experimental data. As shown in **Supplementary Table 6**, 8 patients (corresponding to 8 neurons) were involved in the Vim recordings, from which five patients received DBS across all the frequencies in {5, 10, 20, 30, 50, 100, and 200Hz}. The 200Hz DBS was not applied to three patients, and the 10Hz DBS was not applied to one of these patients (**Supplementary Table 6 – Vim**). To perform the leave-one-out validation, we left out the data from one patient, and fitted the model based on the remaining data; we repeated this procedure for each of the 8 patients. The rate model parameters of the leave-one-out fits were shown in **Supplementary Table 7**, and one can see that estimated parameters are not far from those obtained by incorporating full data (as also shown in **Table 2 – Experimental Data – Vim**). For each rate model parameter, the error percentage of a leave-one-out fit was computed as |“leave-one-out fit” – “full data fit”| / “full data fit”. The mean error percentage across all parameters and all leave-one-out fits was 8.5% (and < 20% across all parameters); confirming that the rate model parameter fitting is robust to the variability caused by removing spikes recorded from an individual patient.

**Supplementary Table 1.** **–** Proportions of excitatory and inhibitory synapses

|  | number of excitatory synapses | number of inhibitory synapses |
| --- | --- | --- |
| **STN** | 150 (30%) | 350 (70%) |
| **SNr** | 50 (10%) | 450 (90%) |
| **Vim / Rt** | 450 (90%) | 50 (10%) |

**Supplementary Table 2.** **–** Weights and time constants of excitatory and inhibitory synaptic currents in 𝐼_𝑠𝑦𝑛_

| | $w_{exc}$ | $w_{inh}$ | $\tau_{exc}$ (ms) | $\tau_{inh}$ (ms) |
| --- | --- | --- | --- | --- |
| **STN** | 1.2 | 1 | 3 | 5 |
| **SNr** | 6 | 4 | 3 | 10 |
| **Vim** | 37.5 | 90 | 5 | 8.5 |
| **Rt** | 4.37 | 11.4 | 5 | 8.5 |

**Supplementary Table 3.** **–** the three types of excitatory synapses

|  | facilitation |  |  | depression |  |  | pseudo linear |  |  |
| --- | --- | --- | --- | --- | --- | --- | --- | --- | --- |
| | $\tau_D$ (ms) | $\tau_F$ (ms) | $U$ | $\tau_D$ (ms) | $\tau_F$ (ms) | $U$ | $\tau_D$ (ms) | $\tau_F$ (ms) | $U$ |
| **STN, SNr,**<br>**Vim / Rt** | 138 | 670 | 0.09 | 671 | 17 | 0.5 | 329 | 326 | 0.29 |

*(B). The ratio of excitatory synaptic types*

|  | facilitation | depression | pseudo linear |
| --- | --- | --- | --- |
| **STN** | 0.1 | 0.6 | 0.3 |
| **SNr** | 0.3 | 0.4 | 0.3 |
| **Vim** | 0.5 | 0.3 | 0.2 |
| **Rt** | 0.5 | 0.3 | 0.2 |

**Supplementary Table 4.** **–** the three types of inhibitory synapses

|  | facilitation |  |  | depression |  |  | pseudo linear |  |  |
| --- | --- | --- | --- | --- | --- | --- | --- | --- | --- |
| | $\tau_D$ (ms) | $\tau_F$ (ms) | $U$ | $\tau_D$ (ms) | $\tau_F$ (ms) | $U$ | $\tau_D$ (ms) | $\tau_F$ (ms) | $U$ |
| **STN, SNr,**<br>**Vim / Rt** | 45 | 376 | 0.016 | 706 | 21 | 0.25 | 144 | 62 | 0.29 |

*(B). The ratio of inhibitory synaptic types*

|  | facilitation | depression | pseudo linear |
| --- | --- | --- | --- |
| **STN** | 0.4 | 0.3 | 0.3 |
| **SNr** | 0.3 | 0.4 | 0.3 |
| **Vim** | 0.3 | 0.4 | 0.3 |
| **Rt** | 0.3 | 0.4 | 0.3 |

**Supplementary Table 5.** **–** Parameters of the background noise current 𝐼_𝑛𝑜𝑖𝑠𝑒_ (in pA) in the LIF model ^1^

| | mean ($\mu$) | st. dev. ($\sigma$) |
| --- | --- | --- |
| **STN** | 32 | 11 |
| **SNr** | 55 | 10 |
| **Vim** | 30 | 45 |
| **Rt** | 12 | 10 |

**Supplementary Table 6.** **–** Number of single-unit recordings in each nucleus for each DBS frequency ^1^

|  | **5Hz** | **10Hz** | **20Hz** | **30Hz** | **50Hz** | **100Hz** | **200Hz** |
| --- | --- | --- | --- | --- | --- | --- | --- |
| **Vim** | 8 | 7 | 8 | 8 | 8 | 8 | 5 |
| **STN** | 6 | 6 | 6 | 6 | 6 | 6 | N/A |
| **SNr** | 5 | 5 | 5 | 5 | 5 | N/A | N/A |
| **Rt** | 7 | 7 | 7 | 7 | 7 | 5 | N/A |

**Supplementary Table 7.** **–** Rate model parameters with leave-one-out fits

| | $\tau_{opt}$ | $r_{b,opt}$ | $c_{opt}$ | $s_{opt}$ | $k_{opt}$ |
| --- | --- | --- | --- | --- | --- |
| full data fit | 45.1 | 13.4 | 687 | $5.81 \times 10^{-2}$ | 548 |
| leave out<br>patient #1 | 43.8 | 15.3 | 744 | $5.87 \times 10^{-2}$ | 545 |
| leave out<br>patient #2 | 47.2 | 15 | 704 | $5.64 \times 10^{-2}$ | 543 |
| leave out<br>patient #3 | 46.3 | 13.1 | 696 | $5.71 \times 10^{-2}$ | 560 |
| leave out<br>patient #4 | 45.1 | 10.5 | 623 | $8.95 \times 10^{-2}$ | 533 |
| leave out<br>patient #5 | 45.5 | 10.0 | 696 | $6.26 \times 10^{-2}$ | 567 |
| leave out<br>patient #6 | 44.0 | 17.7 | 664 | $5.31 \times 10^{-2}$ | 559 |
| leave out<br>patient #7 | 42.2 | 12.3 | 709 | $7.15 \times 10^{-2}$ | 536 |
| leave out<br>patient #8 | 45.7 | 15.7 | 645 | $7.84 \times 10^{-2}$ | 540 |
| mean error<br>percentage | 2.7% | 16.6% | 4.4% | 16.8% | 1.9% |

**Supplementary Figure 1.**
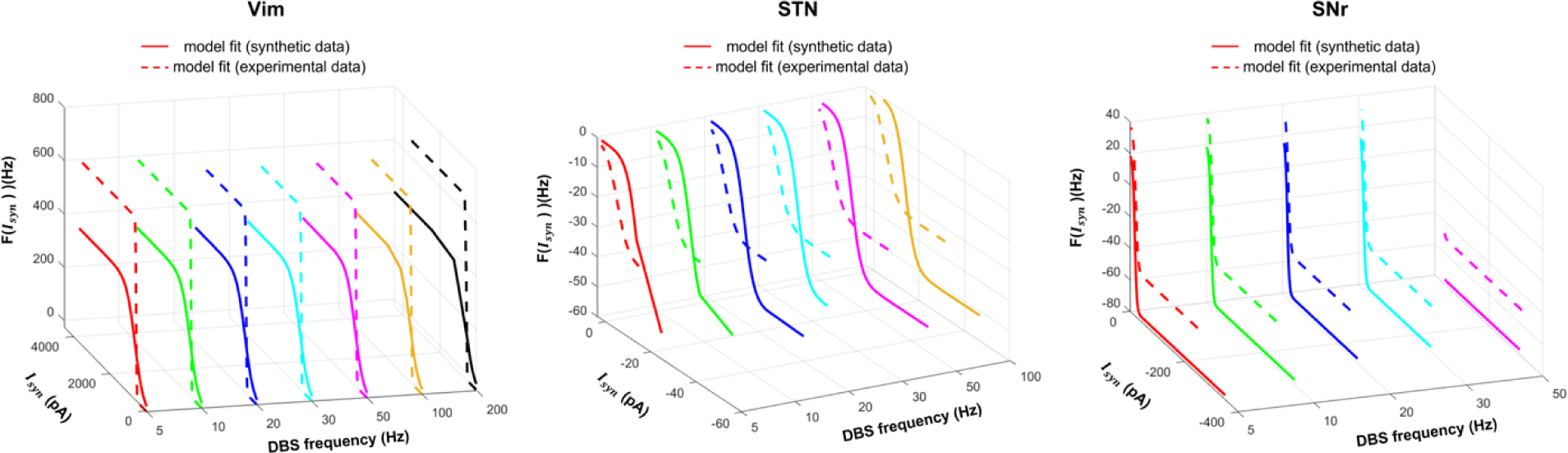
Sigmoid nonlinearity in response to the input synaptic current. The firing rate variability caused by the input synaptic current (I_syn_) was modeled by the sigmoid transfer function (𝐹(𝐼_𝑠𝑦𝑛_), Equation (6)). The unit of 𝐼_𝑠𝑦𝑛_ is picoampere (pA) ^51, 52^. We presented the model fitted transfer functions in both synthetic and experimental data; the synthetic data were from the spiking model in Milosevic et al. (2021) ^1^, and the experimental data were single-unit microelectrode recordings. We showed the transfer functions across all Deep Brain Stimulation (DBS) frequencies, in each of the three basal ganglia and thalamic nucleus: ventral intermediate nucleus (Vim), subthalamic nucleus (STN), and substantia nigra pars reticulata (SNr).

**Supplementary Figure 2.**
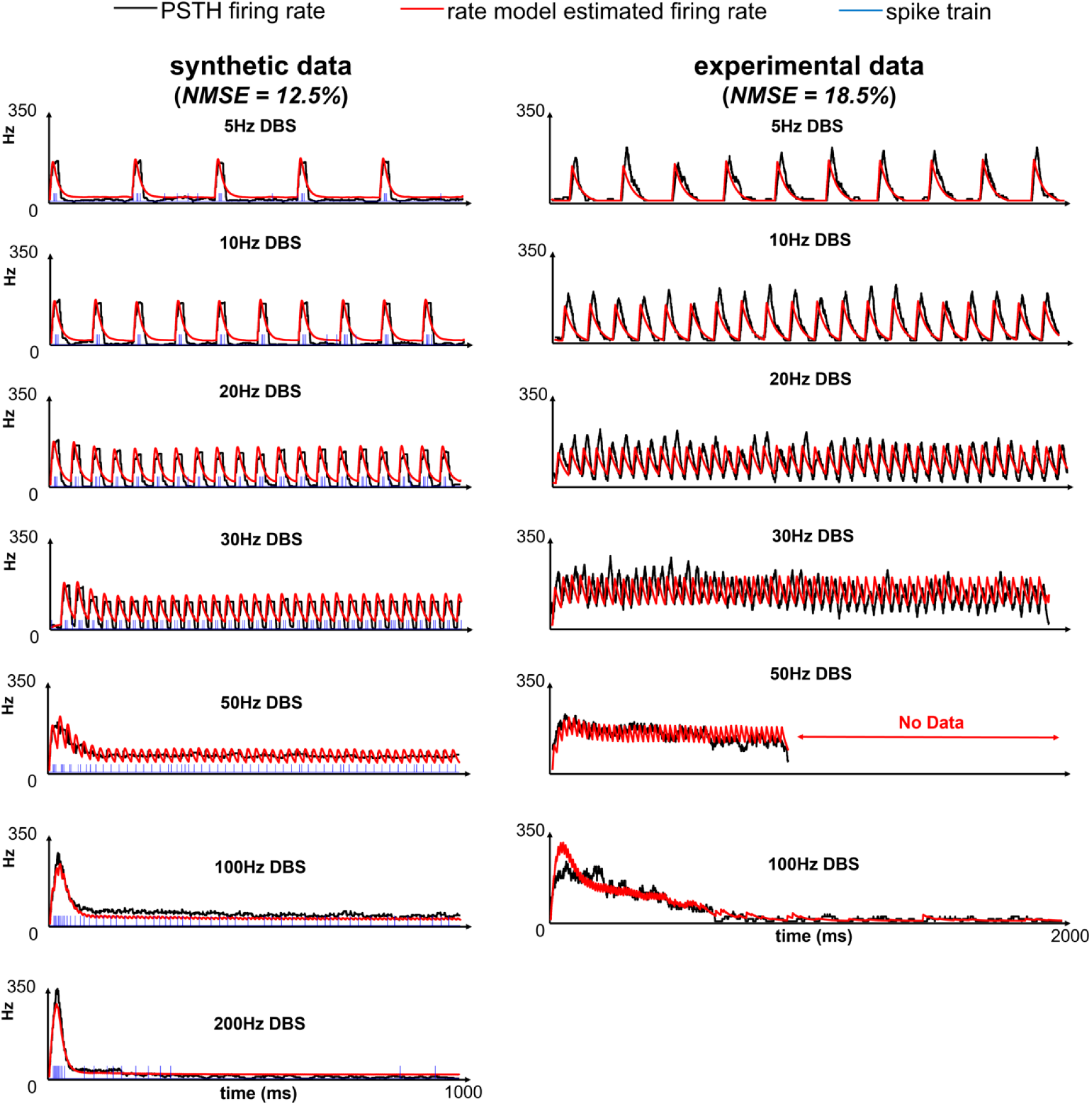
Rate model results for Rt. The rate model was fitted to both synthetic and experimental data from reticular thalamic nucleus (Rt) receiving Deep Brain Stimulation (DBS). The synthetic data were the simulated membrane potentials from the leaky integrate-and-fire (LIF) model established in Milosevic et al. (2021) ^1^, and the experimental data were the single-unit microelectrode recordings. We compared the firing rate computed by peristimulus time histogram (PSTH), firing rate predicted by our rate model and the spike timings from one spike train (for synthetic data only). DBS stimuli with varying pulse frequencies were delivered, and the normalized mean squared error (NMSE) was computed based on the concatenated signal from all DBS frequencies.

**Supplementary Figure 3.**
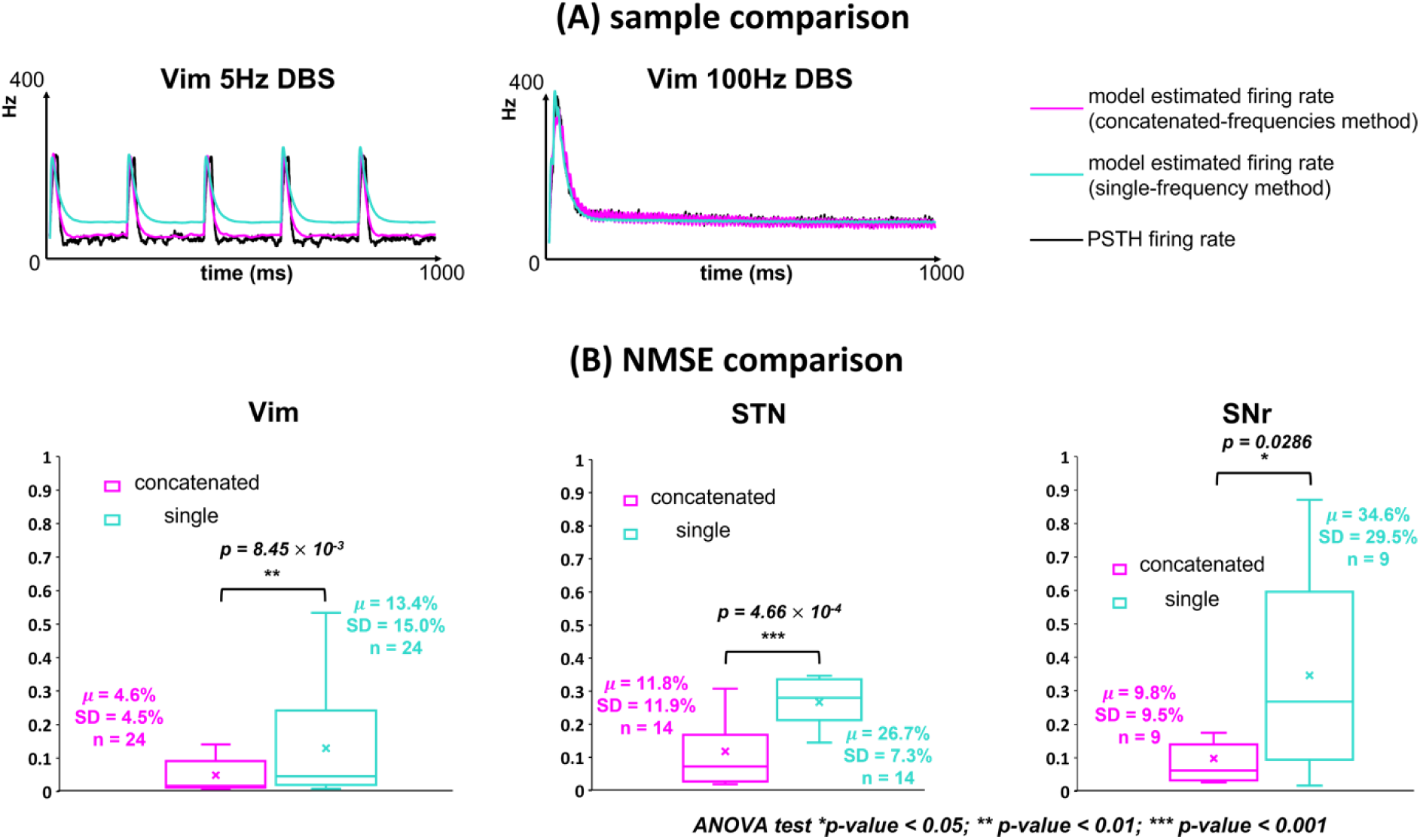
Comparison of two optimization methods in the rate model. “Single-frequency” and “concatenated-frequencies” optimization methods were compared for three basal ganglia and thalamic nuclei: ventral intermediate nucleus (Vim), subthalamic nucleus (STN), and substantia nigra pars reticulata (SNr). (A) The two plots compared the prediction results of the two methods on Vim with varying DBS frequencies. We compared the firing rate computed by peristimulus time histogram (PSTH), and the firing rate predicted by our rate model with each of the two optimization methods. PSTH firing rate was computed based on the synthetic data, i.e., the simulated spike trains of the spiking model in Milosevic et al. (2021) ^1^. For Vim, data from 100Hz DBS was used to train model parameters in the single-frequency method. (B) Normalized mean squared error (NMSE) of the model prediction of multiple DBS frequencies (see text for details) was calculated based on the reference PSTH firing rate. The NMSE results were presented with the box-whisker plot; “concatenated” and “single” mean concatenated-frequencies method and single-frequency method, respectively. “ 𝜇 ” represents the mean value, “SD” represents the standard deviation, and “n” represents the number of samples. ANOVA represents “the one-way analysis of variance test”.

**Supplementary Figure 4.**
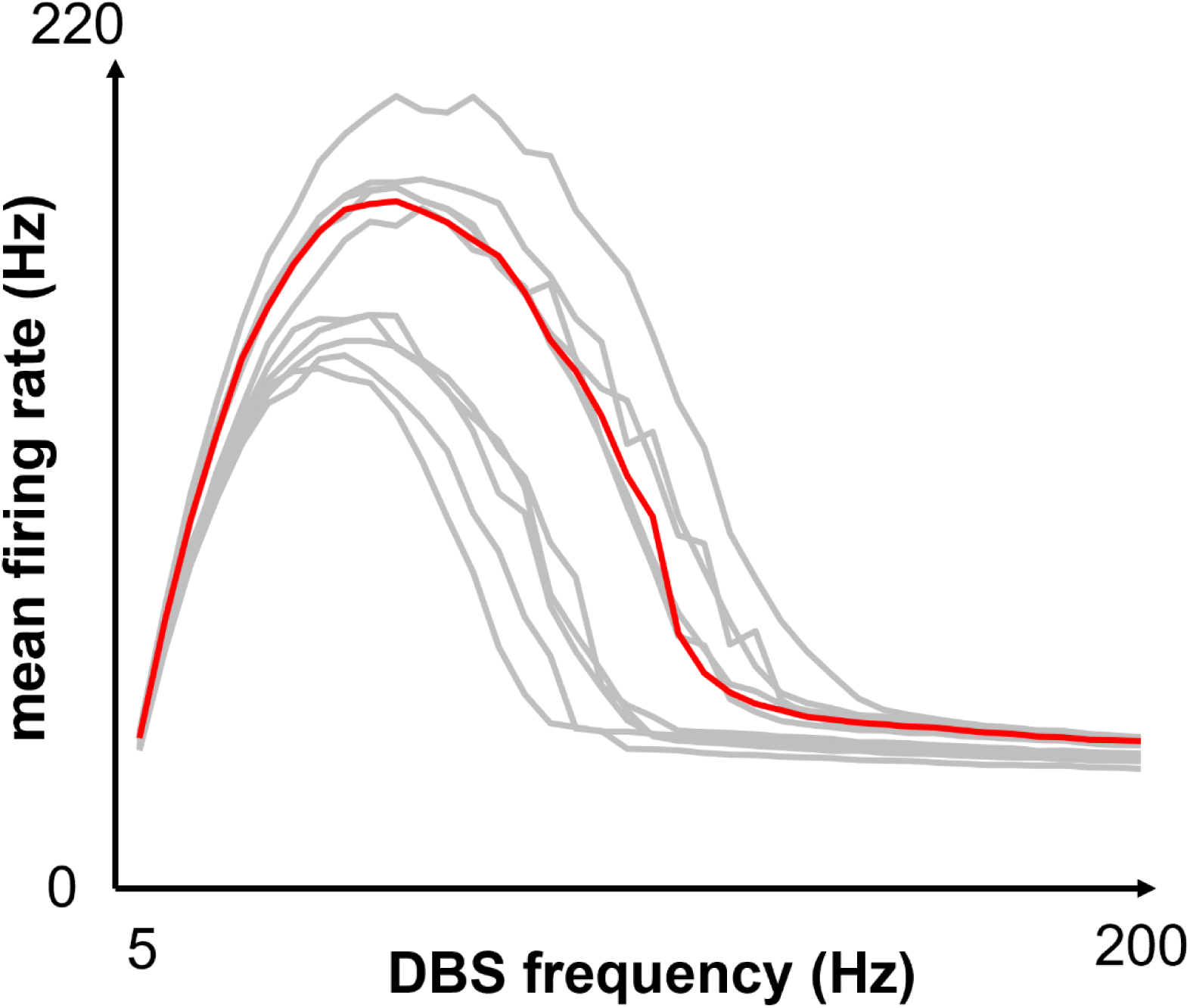
Sensitivity Analysis of the Input Parameters for fitting the rate model to Vim Experimental Data. Simulations were performed at 5 Hz steps in the DBS frequency range [5, 200] Hz. The red trace indicates the synaptic input parameters used in this study, and the gray traces represent the 10 sets of randomized synaptic input parameters in **Supplementary Tables 1 – 4**. We showed the mean firing rate during each simulation.

## Funding

This project was supported by Milad Lankarany’s NSERC Discovery Grant (RGPIN-2020-05868). Leon A Steiner received grant support from Deutsche Forschungsgemeinschaft (DFG, German Research Fundation) – Project-ID 424778381 – TRR 295. Leon A Steiner received additional funding through the Junior Clinician Scientist Program of the Berlin Institute of Health and though a postdoc scholarship provided by the German Academic Exchange Service - DAAD.

## Author Contributions

Yupeng Tian and Milad Lankarany were responsible for the conception and design of the work. Leon A Steiner, Suneil K Kalia, Mojgan Hodaie, Andres M Lozano, William D Hutchison and Luka Milosevic were responsible for the acquisition of data. Yupeng Tian and Matthew J. H. Murphy were responsible for analyzing the data. Yupeng Tian, Milos R. Popovic, and Milad Lankarany were responsible for interpreting the data. Yupeng Tian and Milad Lankarany were responsible for drafting the work. All authors are responsible for revising the work critically for important intellectual content. All authors approved the final version of the manuscript, and agree to be accountable for all aspects of the work in ensuring that questions related to the accuracy or integrity of any part of the work are appropriately investigated and resolved. All persons designated as authors qualify for authorship, and all those who qualify for authorship are listed.

## Conflict of Interest

All authors declare no conflict of interest.

## Notes

### Competing Interest Statement

The authors have declared no competing interest.

### Summary of Updates

add model validation and more statistical information

